# Human cytomegalovirus UL7, miR-US5-1 and miR-UL112-3p inactivation of FOXO3a protects CD34^+^ hematopoietic progenitor cells from apoptosis

**DOI:** 10.1101/2020.09.28.317859

**Authors:** Meaghan H. Hancock, Lindsey B. Crawford, Wilma Perez, Hillary M. Struthers, Jennifer Mitchell, Patrizia Caposio

## Abstract

Human cytomegalovirus (HCMV) infection of myeloid-lineage cells, such as CD34^+^ hematopoietic progenitor cells (HPCs) or monocytes results in the upregulation of anti-apoptotic cellular proteins that protect the newly infected cells from programmed cell death. The mechanisms used by HCMV to regulate pro-apoptotic cellular proteins upon infection of CD34^+^ HPCs has not been fully explored. Here we show that HCMV utilizes pUL7, a secreted protein that signals through the FLT3 receptor, and miR-US5-1 and miR-UL112-3p to reduce the abundance and activity of the pro-apoptotic transcription factor FOXO3a at early times after infection of CD34^+^ HPCs. Regulation of FOXO3a by pUL7, miR-US5-1 and miR-UL112 results in reduced expression of the pro-apoptotic *BCL2L11* transcript and protection of CD34+ HPCs from virus-induced apoptosis. These data highlight the importance of both viral proteins and miRNAs in protecting CD34^+^ HPCs from apoptosis at early times post-infection, allowing for the establishment of latency and maintenance of viral genome-containing cells.

**Importance:** Human cytomegalovirus (HCMV) causes serious disease in immunocompromised individuals and is a significant problem during transplantation. The virus can establish a latent infection in CD34^+^ hematopoietic progenitor cells (HPCs) and periodically reactivate to cause disease in the absence of an intact immune system. What viral gene products are required for successful establishment of latency are still not fully understood. Here we show that both a viral protein and viral miRNAs are required to prevent apoptosis after infection of CD34^+^ HPCs. HCMV pUL7 and miRNAs miR-US5-1 and miR-UL112-3p act to limit the expression and activation of the transcription factor FOXO3a which in turn reduces expression of pro-apoptotic gene *BCL2L11* and prevents virus-induced apoptosis in CD34^+^ HPCs.

## Introduction

Human Cytomegalovirus (HCMV) is prevalent in the majority of the world’s population and results in a lifelong persistence with periods of latency and reactivation. Primary infection in healthy individuals is typically sub-clinical and is controlled by the host immune response. However, HCMV infection of immune compromised hematopoietic stem cell (HSCT) or solid organ (SOT) transplant recipients remains a major cause of life-threatening disease even with the development of antiviral therapy (1-3). The main source of latent HCMV are hematopoietic progenitor cells (HPCs) and myeloid lineage cells (4-6). HCMV expresses a limited set of genes during latency including US28, transcriptional variants from the UL133-138 region, LUNA, a subset of viral miRNAs and the β2.7 long non-coding RNA (lncRNA). Viral reactivation from latency is tightly linked to myeloid differentiation that results in the expression of all the genes necessary to produce infectious virus in macrophages (7, 8). The HCMV latency genes function through modulation of cellular processes necessary for maintenance of the viral genome in the cell. The triggers for viral reactivation and the viral and cellular genes that mediate this process are largely unknown.

HCMV infection of cells results in the activation of pro-apoptotic cell death pathways that are detrimental to viral replication. HCMV encodes a number of anti-apoptotic proteins that function during lytic infection to block this cellular response. These HCMV anti-apoptotic proteins include pUL36 (vICA) that interacts with procaspase-8 to inhibit proteolytic processing (9), pUL37 (vMIA) that sequesters proapoptotic BAX at the outer mitochondrial membrane and prevents cytochrome c release (10), and the pUL38 gene product that blocks proteolysis of two key apoptotic enzymes, caspase-3 and poly(ADP-ribose) polymerase (11). In addition to these proteins, the HCMV immediate early protein IE2 was observed to upregulate the protease-deficient procaspase-8 homologue, c-FLIP, that decreases the activities of caspase 3 and 8 (12). Also, HCMV lncRNA β2.7 was shown to interact with the mitochondrial enzyme complex I to stabilize mitochondrial membrane potential and prevent apoptotic death of HCMV-infected neuronal cells (13). Several HCMV miRNAs have also been described that protect cells from apoptosis during lytic infection. HCMV miR-UL36-5p inhibits apoptosis via downregulation of adenine nucleotide translocator 3, an adenine nucleotide transporter responsible for translocating ADP and ATP across the mitochondrial membrane (14). Additionally, miR-UL148D protects cells from apoptosis induced by overexpression of IEX-1 (15). Finally, Kim *et al*. reported *FAS* as a target of miR-UL36-3p, miR-US5-1, and miR-US5-2-3p, *Fas associated protein with death domain (FADD)* as a target of miR-US5-1, *Caspase 3* as a target of miR-US25-2-3p, miR-112-5p, and miR-UL22A-5p, *Caspase 2* as a target of miR-US4-5p and *Capase 7* as a target of miR-UL22A-3p and miR-US4-3p (16).

In non-permissive cells (CD34^+^ HPCs), or in cells with a protracted life cycle (monocytes), early survival of HCMV-infected cells is achieved by virus-induced regulation of anti-apoptotic cellular factors such as myeloid cell leukemia (MCL)-1 and B cell lymphoma-2 (Bcl-2) (reviewed in (17)). In monocytes, the binding of HCMV gB to epidermal growth factor receptor (EGFR), as well as the binding of the viral pentameric complex to integrins drives signaling through PI3K and mTOR kinase that leads to the up-regulation of MCL-1 (18). The virus, after 48 hours of infection, shifts from MCL-1 to Bcl-2 as the primary anti-apoptotic tool and the upregulation of Bcl-2 is mediated by integrin signaling events following initial viral binding (19). In CD34^+^ HPCs, Reeves *et al*. showed that MCL-1 is upregulated via gB stimulation of MAPK signaling in the absence of *de novo* viral gene expression (20). Furthermore, the protective phenotype driven by the virus-induced ERK-MAPK signaling correlates with the downregulation of the pro-apoptotic proteins PUMA and BIM. At the same time, ERK-MAPK signaling phosphorylates the transcription factor ELK-1 that is required for MCL-1 expression and cell survival (21).

Expression of cellular pro- and anti-apoptotic genes is carefully regulated to allow for the timely elimination of transformed and virus-infected cells. The mammalian Forkhead Box O (FOXO) family of transcription factors, including FOXO1, FOXO3a, FOXO4, and FOXO6, are implicated in a wide variety of physiologic processes such as cell cycle arrest, cellular differentiation, DNA repair and cell death (reviewed in (22, 23)). The FOXO family of transcription factors promote apoptosis by mitochondrial-dependent and -independent mechanisms, including mediating expression of the Bcl-2-like protein 11 (BIM), a pro-apoptotic Bcl-2 family protein (24). FOXO proteins are normally present in an active state in the cell’s nucleus. Upon growth factor stimulation they are phosphorylated by different serine/threonine cellular kinases triggering inactivation and export of FOXOs from the nucleus to the cytoplasm (25) (Reviewed in (26)).

HCMV UL7 is part of the RL11 gene family and is dispensable for lytic viral replication (27). Our group and others have shown that UL7 is a transmembrane glycoprotein that is secreted from infected cells (27-29). Among clinical and lab adapted HCMV strains, UL7 sequence in very well conserved (97 to 100% intragenotype conservation and 83 to 93% intergenotype conservation), suggesting a crucial role in viral replication in the host (30). Our group recently demonstrated that HCMV pUL7 is a ligand for the cytokine receptor Fms-like tyrosine kinase 3 (Flt-3R) (31). Signaling through the Flt-3R is critical for normal development of hematopoietic stem and progenitor cells (32). We observed that pUL7 binding to Flt-3R induces activation of the downstream PI3K/AKT and MAPK/ERK signaling pathways. Importantly, we have shown that UL7 protein induces both CD34^+^ HPC as well as monocyte differentiation *in vitro* and *in vivo* and thus functions as a hematopoietic differentiation factor (31). Although UL7 is non-essential for lytic replication, HCMV UL7 mutants fail to reactivate from latency in CD34^+^ HPC (31).

In the current study we observed that pUL7 signaling *via* Flt-3R promotes a rapid phosphorylation of FOXO3a specifically through the MAPK pathway. The phosphorylation of FOXO3a results in nuclear-to-cytoplasmic translocation and inactivation of the transcription factor as demonstrated by the downregulation of the FOXO3a target gene *BCL2L11*. Additionally, we show that HCMV miR-US5-1 and miR-UL112-3p directly reduce FOXO3a transcript and protein levels, resulting in reduced *BCL2L11* mRNA expression, indicating that the virus utilizes multiple mechanisms to modulate FOXO3a activity. Finally, we observed that UL7, miR-US5-1, and miR-UL112-3p are expressed in the early stages of HCMV infection in CD34^+^ HPCs and act to reduce FOXO3a activity to promote survival of infected hematopoietic progenitor cells.

## Results

### pUL7 signaling promotes phosphorylation of FOXO3a via the MAPK pathway

We recently demonstrated that pUL7 binds and signals through the Flt-3R (31). Flt-3 ligand (Flt-3L) induction of AKT/PKB activation was reported to lead to phosphorylation and inactivation of FOXO3a (33) but since MAPK/ERK signaling can also lead to FOXO3a phosphorylation (34), we first determined if pUL7 was able to promote phosphorylation of FOXO3a and which signaling pathway was involved in this process. Since few primary cell types express the Flt-3R, we established Flt-3L responsive cells by retroviral gene transfer the human *Flt-3R* gene into telomerized human fibroblasts (THF). When THF-Flt-3R cells were stimulated by pUL7 or the control Flt-3L, FOXO3a was rapidly phosphorylated in a time dependent manner (Figure 1A) and the response was dependent on the Flt-3R as demonstrated by the treatment with the Flt-3R inhibitor AC220 (Figure 1B). To determine which Flt-3R downstream signaling pathway was involved in FOXO3a phosphorylation we stimulated the cells in the presence of a PI3K inhibitor (LY294002), or a MEK inhibitor (PD98059), and we used as a control a S6K inhibitor (LY303511) to rule out off target effects. As previously reported, phosphorylation of FOXO3a by Flt-3L was PI3K-dependent as addition of LY294002 decreased the level of FOXO3a phosphorylation, while pUL7 resulted in phosphorylation of FOXO3a through the MAPK pathway. Indeed, neither LY294002 nor LY303511 had any effect on pUL7-mediated FOXO3a phosphorylation, only the MAP kinase inhibitor PD98059 was able to prevent pUL7-induced FOXO3a phosphorylation (Figure 1C).

**Figure 1.**
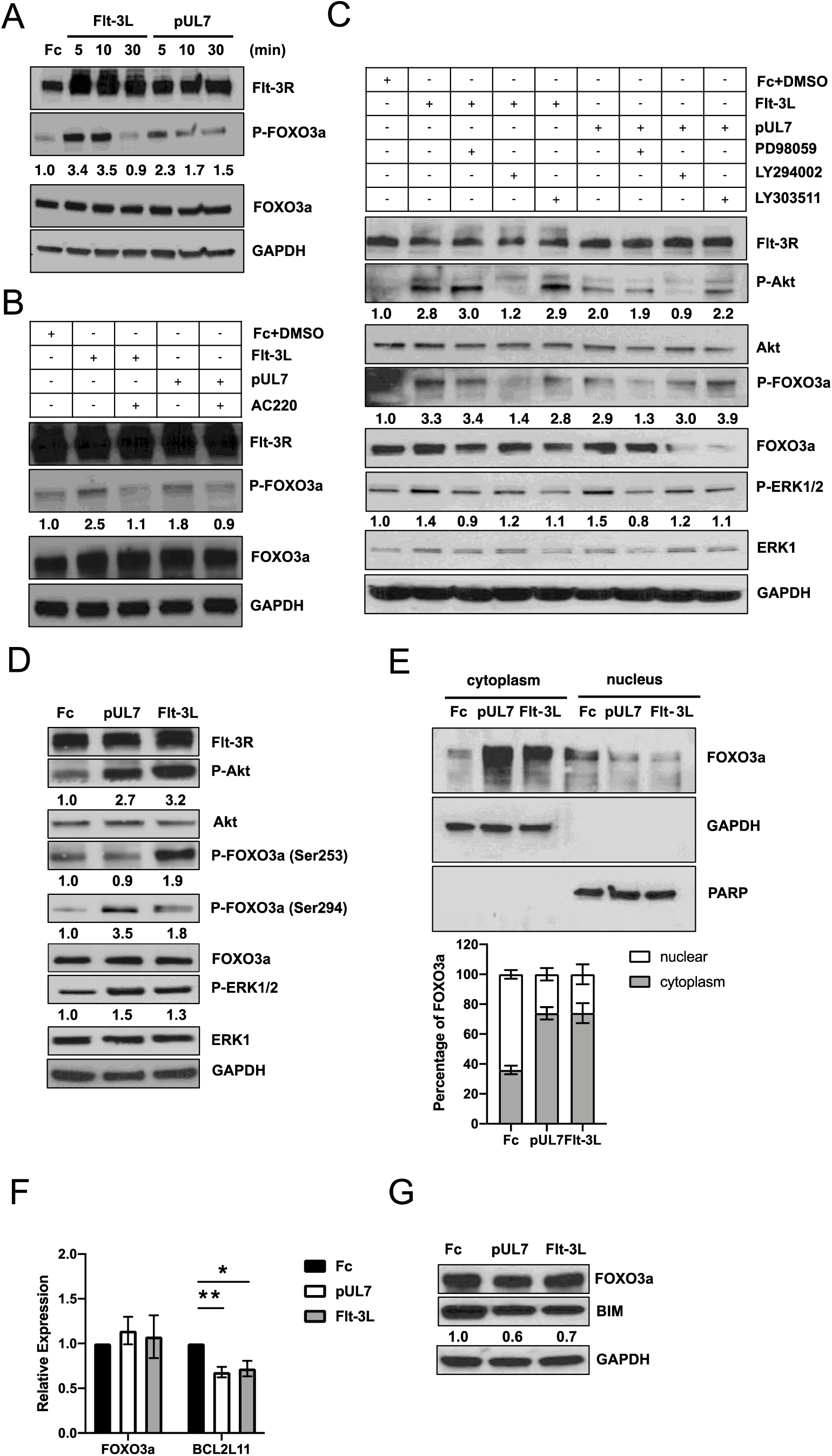
pUL7 promotes FOXO3a phosphorylation via MAPK pathway. A) Protein lysates were generated and immunoblotted for phosphorylation of FOXO3a from serum-starved THF-Flt-3R stimulated with 50 ng/ml of Fc, pUL7, or Flt-3L for the indicated time. B) THF-Flt-3R were pre-treated with AC220 (100 nM) for 1 h and then stimulated for 10 min with Fc + DMSO, pUL7, or Flt-3L. Protein lysates were generated and immunoblotted for phosphorylation of FOXO3a (Ser253/Se294). C) THF-Flt-3R were pre-treated with PD98059 (50 μM), LY294002 (20 μM), LY303511 (25 μM) for 1 h and then stimulated for 10 min. Protein lysates were generated and immunoblotted for phosphorylation of Akt (Thr308), ERK1/2 (Thr202/Tyr204), FOXO3a. D) Protein lysates were generated from THF-Flt-3R stimulated for 10 min and immunoblotted for phosphorylation of Akt (Thr308), ERK1/2 (Thr202/Tyr204), FOXO3a (Ser253), and FOXO3a (Ser 294). Equal loading was confirmed by Flt-3R, Akt, ERK1, FOXO3a, and GAPDH antibody staining. Numbers under the immunoblots indicate relative phosphorylation of FOXO3a, Akt, or ERK1/2 normalized to the amount of total FOXO3a, Akt or ERK1 and compared to the value of the Fc control. E) Nuclear versus cytoplasmic lysates were generated from THF-Flt-3R stimulated for 10 min and immunoblotted for total FOXO3a. GAPDH and PARP were used as cytosolic and nuclear markers, respectively. Relative intensity was quantified by densitometry and normalized to the loading controls. The percentage of cytoplasmic and nuclear FOXO3a was determined as follow: cytoplasmic FOXO3a density/(cytoplasmic+ nuclear FOXO3a density) x100 and nuclear FOXO3a density/(cytoplasmic+ nuclear FOXO3a density) x100, respectively. Results are representative of three independent experiments. F) Serum-starved THF-Flt-3R cells were stimulated for 24 hours with Fc, pUL7, or Flt-3L. RNA was isolated using Trizol and qRT-PCR for *FOXO3a* and the downstream target *BCL2L11* was performed. Values are means±standard error of the means (SEM) (error bars) from three independent experiments. Statistical significance was determined by unpaired Student’s t-test (*, *P<0*.*05*, **; *P<0*.*005*; ***; *P<0*.*0005*).

To further confirm this observation, we used specific phospho-FOXO3a antibodies that recognize serine residues phosphorylated by Akt (Ser253) or ERK1/2 (Ser294) (26). As shown in Figure 1D, stimulation with Flt-3L induced rapid phosphorylation of FOXO3a on Ser253, while pUL7 stimulation resulted in FOXO3a phosphorylation on Ser294. Overall, FOXO3a levels were lower in the cytoplasm of the control (35.99% ± 2.89) compared to the Flt-3L (74.05% ± 6.66) and pUL7 (73.91% ± 4.12) stimulated cells (Figure 1E), further supporting the observation that FOXO3a is inactivated by pUL7.

Based on the observed phosphorylation and re-localization of FOXO3a to the cytoplasm after pUL7 treatment, we next tested whether transcription of FOXO3a target gene *BCL2L11* was also affected. THF-Flt-3R cells were placed in low serum media, and then treated with pUL7 or Flt-3L for 24 hours. As shown in Figure 1F, we observed a significant reduction in *BCL2L11* expression after pUL7 (p=0.0054) or Flt-3L (p=0.0325) stimulation. The reduction in *BCL2L11* mRNA also affected protein levels (Figure 1G). Finally, stimulation with pUL7 or Flt-3L did not affect FOXO3a mRNA or protein expression confirming that pUL7 regulates FOXO3a via post-translational modifications without affecting mRNA or total protein levels (Figure 1F and G).

### pUL7 promotes cytoplasmic translocation of FOXO3a and downregulation of *BCL2L11* in myeloid cells

To validate our findings in myeloid cells, we stimulated Flt-3R expressing bone marrow lymphoblast RS4;11 cells with pUL7 in the presence or absence of different chemical inhibitors. As previously observed, pUL7-mediated phosphorylation of FOXO3a was inhibited by the MEK inhibitor (PD98059), but not by the PI3K inhibitor (LY294002) or the control S6K inhibitor (LY303511) (Figure 2A). When we examined how FOXO3a phosphorylation affects the distribution of protein between the cytoplasm and nucleus, we found that in Fc treated cells FOXO3a was mostly nuclear at each time point post-treatment. However, upon pUL7 stimulation, FOXO3a translocated into the cytoplasm, with significantly less FOXO3a in the nucleus after 6 hours (p=0.0033, Figure 2B). Next, we analyzed FOXO3a translocation in primary CD34^+^ HPC, and as shown in Figure 2C, FOXO3a protein is more abundant in the cytoplasm of pUL7 stimulated cells compared to the control (p=0.0179). Furthermore, FOXO3a deficiency in the nucleus via pUL7 stimulation decreases the transcript levels of *BCL2L11* in CD34^+^ HPCs (p=0.0278, Figure 2D).

**Figure 2.**
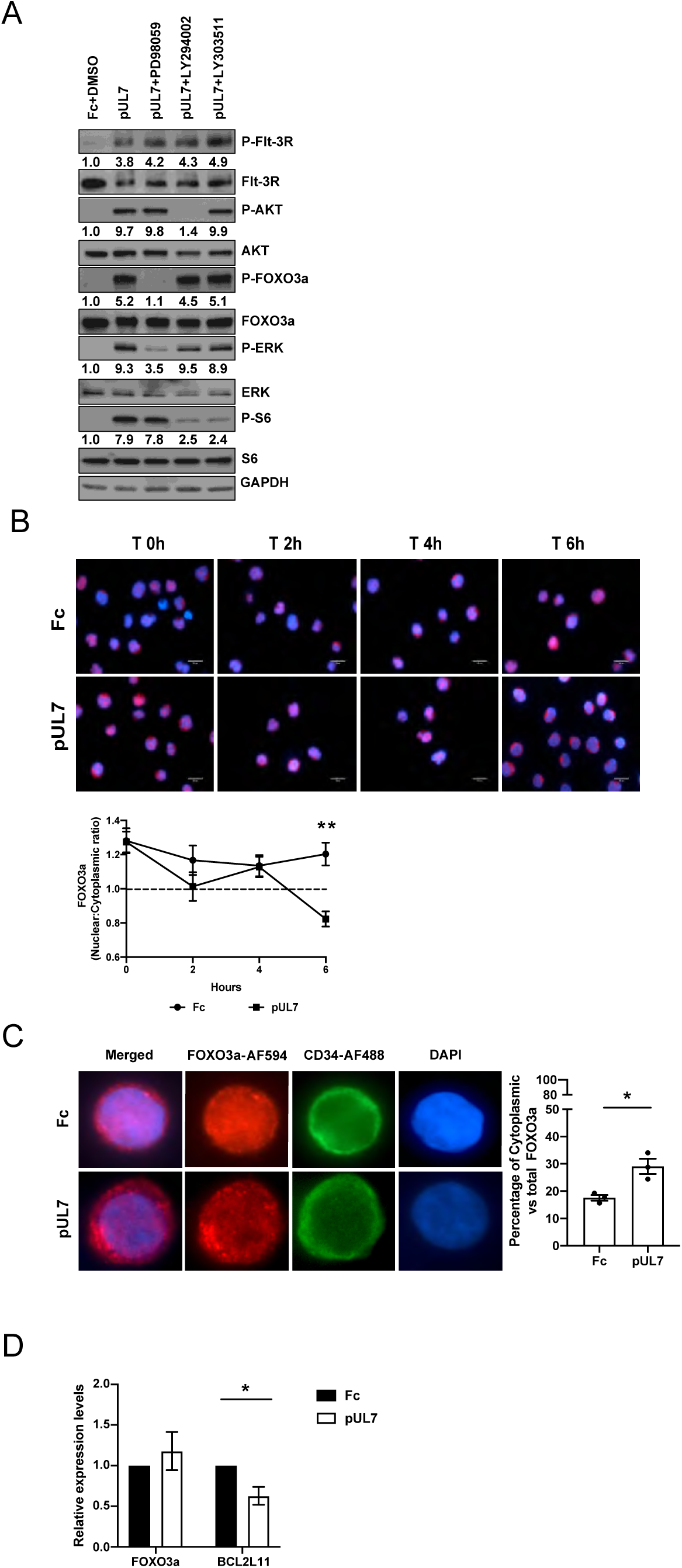
pUL7 induces FOXO3a translocation from the nucleus to the cytoplasm in RS4;11 and CD34^+^ HPC cells. A) RS4;11 cells were serum-starved, pre-treated with PD98059 (50 μM), LY294002 (20 μM), LY303511 (25 μM) for 1 hour and then stimulated for 10 min with Fc+DMSO or pUL7. Protein lysates were generated and immunoblotted for phosphorylation of Flt-3R (Tyr591), Akt (Thr308), ERK1/2 (Thr202/Tyr204), FOXO3a (Ser 294), and S6 (Ser235/236). Equal loading was confirmed by Flt-3R, Akt, ERK1, FOXO3a, S6 and GAPDH antibody staining. Numbers under the immunoblots indicate relative phosphorylation of Flt-3R, Akt, FOXO3a, ERK1/2 or S6 normalized to the amount of total Flt-3R, Akt, FOXO3a, ERK1, and S6 and compared to the value of the Fc control. B) Subcellular localization of FOXO3a in RS4;11 cells stimulated with Fc and pUL7 (50 ng/ml) at the indicated time points (representative images). FOXO3a (red, secondary antibody conjugated to AF594) and nucleus (DAPI, blue). Images were quantified using ImageJ and the ratio of nuclear to cytoplasmic FOXO3a over time is shown. The dotted line indicate transition from higher signal in the nucleus (above) to cytoplasm (below). Average of 30 cells were analyzed per condition. Values are means±standard error of the means (SEM) (error bars) from three independent experiments. Statistical significance was determined by unpaired Student’s t-test (*, *P<0*.*05*, **; *P<0*.*005*; ***; *P<0*.*0005*). C) Bone marrow CD34^+^ HPCs were stimulated for 6 hours with Fc or pUL7 (50 ng/ml) and then stained with anti-FOXO3a antibody and Alexa Fluor AF594conjugated secondary antibody to detect FOXO3a (red), anti-CD34 antibody and Alexa Fluor AF488-conjugated secondary antibody to detect CD34 (green). DAPI was used to counterstain the nucleus (blue). Quantification was done comparing percentage of cytoplasmic to the total FOXO3a. Average of 30 cells were analyzed per condition. Values are means±standard error of the means (SEM) (error bars) from three independent donors. Statistical significance was determined by unpaired Student’s t-test (*, *P<0*.*05*, **; *P<0*.*005*; ***; *P<0*.*0005*). D) Bone marrow CD34^+^ HPCs were stimulated for 6h with Fc or pUL7. RNA was isolated using Trizol and qRT-PCR for *FOXO3a* and the downstream target gene *BCL2L11* was performed. Values are means±standard error of the means (SEM) (error bars) from three independent donors. Statistical significance was determined by unpaired Student’s t-test (*, *P<0*.*05*, **; *P<0*.*005*; ***; *P<0*.*0005*).

Taken together, these data demonstrate that pUL7 plays a pivotal role in the phosphorylation and translocation of FOXO3a from the nucleus to the cytoplasm in progenitors and myeloid cells, leading to inactivation of the transcription factor and reduction in expression of the pro-apoptotic gene *BCL2L11*.

### HCMV miR-US5-1 and miR-UL112 cooperatively target FOXO3a

There are numerous examples of redundancy between herpesvirus proteins and miRNAs (35, 36). For example, FOXO3a is targeted for inactivation by both gammaherpesvirus proteins and miRNAs (37). Thus, we investigated whether FOXO3a could also be a target of HCMV miRNAs. Biochemical and bioinformatic analysis (16) suggested that FOXO3a was a target of HCMV miR-US5-1 and miR-UL112-3p. We cloned the 3’ UTR of FOXO3a or 3’ UTRs containing deletions of the potential miR-US5-1 or miR-UL112-3p target sites into a dual luciferase vector. HEK293T cells were transfected with the luciferase vectors along with negative control miRNA, miR-US5-1 or miR-UL112-3p miRNA mimics. Both miR-US5-1 and miR-UL112-3p target the 3’ UTR of FOXO3a through the identified sites (Figure 3A). The HCMV miRNAs are capable of downregulating FOXO3a transcript (p<0.0001, Figure 3B) and endogenous protein (Figure 3C) after transfection of miRNA mimics into human fibroblasts. Since transfection efficiency in CD34^+^ HPCs is usually low, we generated GFP-expressing adenoviral vectors additionally expressing miR-US5-1, miR-UL112-3p, or and a FOXO3a shRNA to validate our results in progenitor cells. The other advantage of using the adenovirus system is the presence of a GFP expression cassette that allows for sorting of a pure population of transduced cells. We first validated the function of Ad miR-US5-1, Ad miR-UL112-3p, and Ad FOXO3a shRNA into human fibroblasts. Expression of either the two HCMV miRNAs or the control shRNA downregulates *FOXO3a* (Figure 3D) and *BCL2L11* mRNA (Figure 3E) as well as FOXO3a and BIM protein levels (Figure 3F). These results clearly demonstrate that FOXO3 is a target of HCMV miR-US5-1 and UL112-3p.

**Figure 3.**
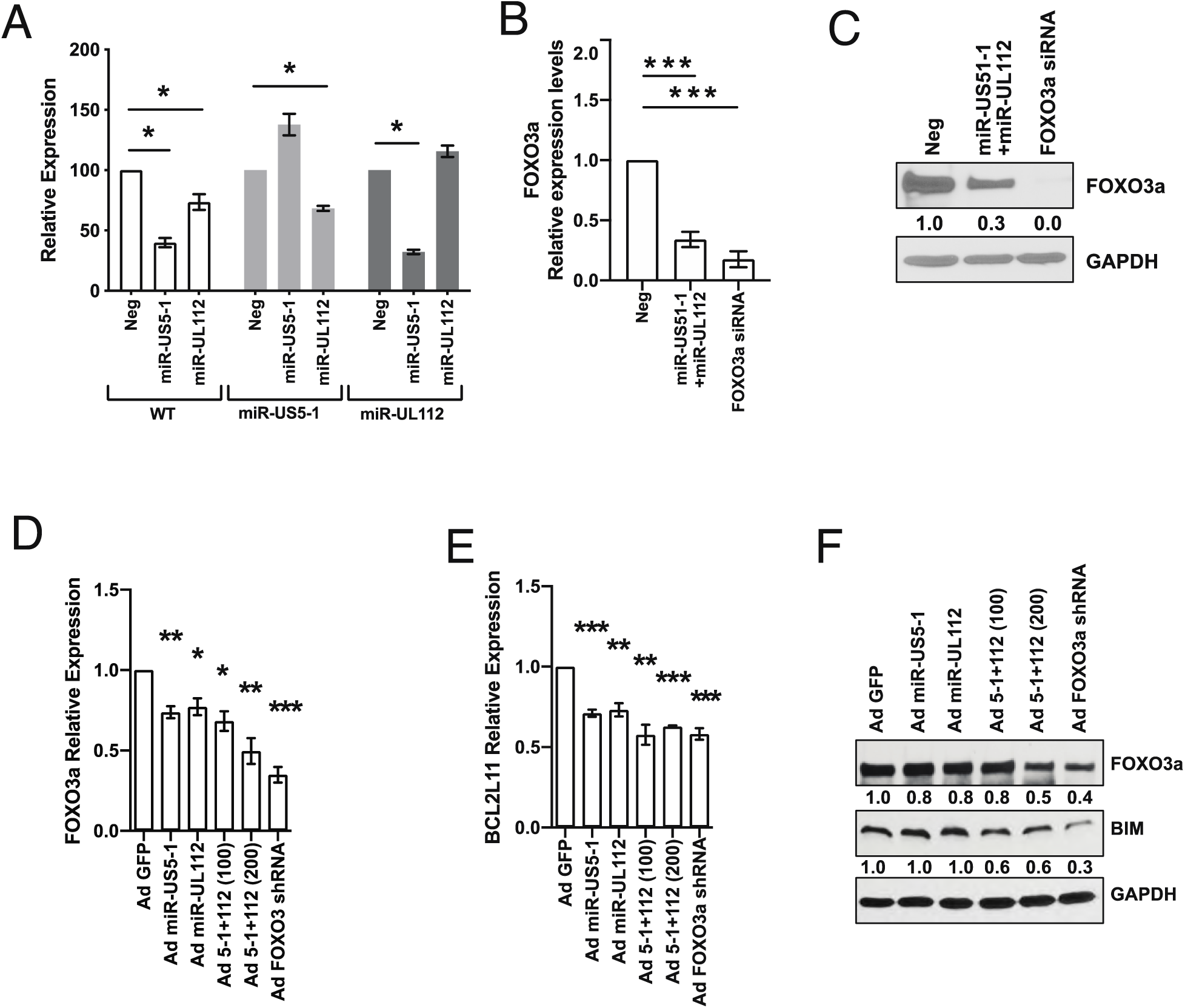
HCMV miR-US5-1 and miR-UL112-3p target FOXO3a protein for downregulation. A) The WT FOXO3a 3’ UTR or 3’ UTRs containing mutations in the miR-US5-1 or miR-UL112 binding sites were cloned into the pSiCheck2 dual luciferase vector. HEK293T cells were transfected with the pSiCheck2 vector along with negative control miRNA, miR-US5-1 or miR-UL112-3p mimic. 24 hours later cells were lysed and luciferase measured. Values are means±standard error of the means (SEM) (error bars) for three independent experiments. Statistical significance was determined by unpaired Student’s t-test (*, *P<0*.*05*, **; *P<0*.*005*; ***; *P<0*.*0005*). B) Normal human dermal fibroblasts (NHDF) were transfected with negative control miRNA, miR-US5-1 and miR-UL112-3p mimics or a FOXO3a siRNA for 48 hours. RNA was isolated using Trizol and qRT-PCR for *FOXO3a* was performed. Values are means±standard error of the means (SEM) (error bars) for three independent experiments. Statistical significance was determined by unpaired Student’s t-test (*, *P<0*.*05*, **; *P<0*.*005*; ***; *P<0*.*0005*). (C) NHDF were transfected as in (B) and protein subjected to immunoblotting. Images were quantified using ImageJ and the ratio of FOXO3a to GAPDH protein, normalized to Neg control. D) NHDF were serum starved and then transduced with the different adenoviral vectors for 48 hours at MOI 200 or 100 when in combination. RNA was isolated using Trizol and qRT-PCR for *FOXO3a* or *BCL2L11* (E) was performed. Values are means±standard error of the means (SEM) (error bars) for three independent experiments. Statistical significance was determined by unpaired Student’s t-test (*, *P<0*.*05*, **; *P<0*.*005*; ***; *P<0*.*0005*). F) NHDF were treated as described in (D) and protein subjected to immunoblotting. Images were quantified using ImageJ and the ratio of FOXO3a and BIM to GAPDH protein, normalized to Ad GFP control.

### Expression of HCMV UL7, miR-US5-1, and miR-UL112-3p protects CD34^+^ HPCs from apoptosis

Next, we additionally generated a GFP-expressing UL7 adenoviral vector. As shown in Figure S1, transduction of human CD34^+^ HPCs with Ad UL7 leads to expression and secretion of UL7. We used each of the adenoviral vectors to transduce CD34^+^ HPCs, then FACS-isolated a pure population of viable CD34^+^ GFP^+^ cells, and determined the effect of viral miRNA or protein expression on *FOXO3a* and *BCL2L11* mRNA levels. As shown in Figure 4A and S2, expression of miR-US5-1, miR-UL112-3p, and the control FOXO3a shRNA downregulates *FOXO3a* (p=0.0001, p=0.0003, p=0.0001) and *BCL2L11* (p=0.0002, p=0.029, p=0.001) mRNA levels. Consistent with its effect only on the phosphorylation status of FOXO3a, expression of UL7 did not have any impact on *FOXO3*a mRNA levels (Figure 4A) but significantly decreased the pro-apoptotic gene *BCL2L11* (p=0.004) (Figure 4B). We then wanted to evaluate the effect of UL7 and HCMV miRNAs on adenoviral-induced apoptosis (38). CD34^+^ HPCs from three independent donors were transduced with the adenoviral vectors and phosphatidylserine translocation, as measured by Annexin V staining on CD34^+^ GFP^+^ HPCs using flow cytometry, was analyzed after 72 hours. As shown in Figure 4C, CD34^+^ HPCs expressing HCMV miR-US5-1 (p=0.02), miR-UL112-3p (p=0.0137), or pUL7 (p=0.0056) were more resistant to apoptosis compared to Ad GFP. These data indicate that inactivation of FOXO3a by HCMV miR-US-5-1, miR-UL112-3p, and UL7 results in decreased levels of *BCL2L11* and ultimately protection from apoptosis.

**Figure 4.**
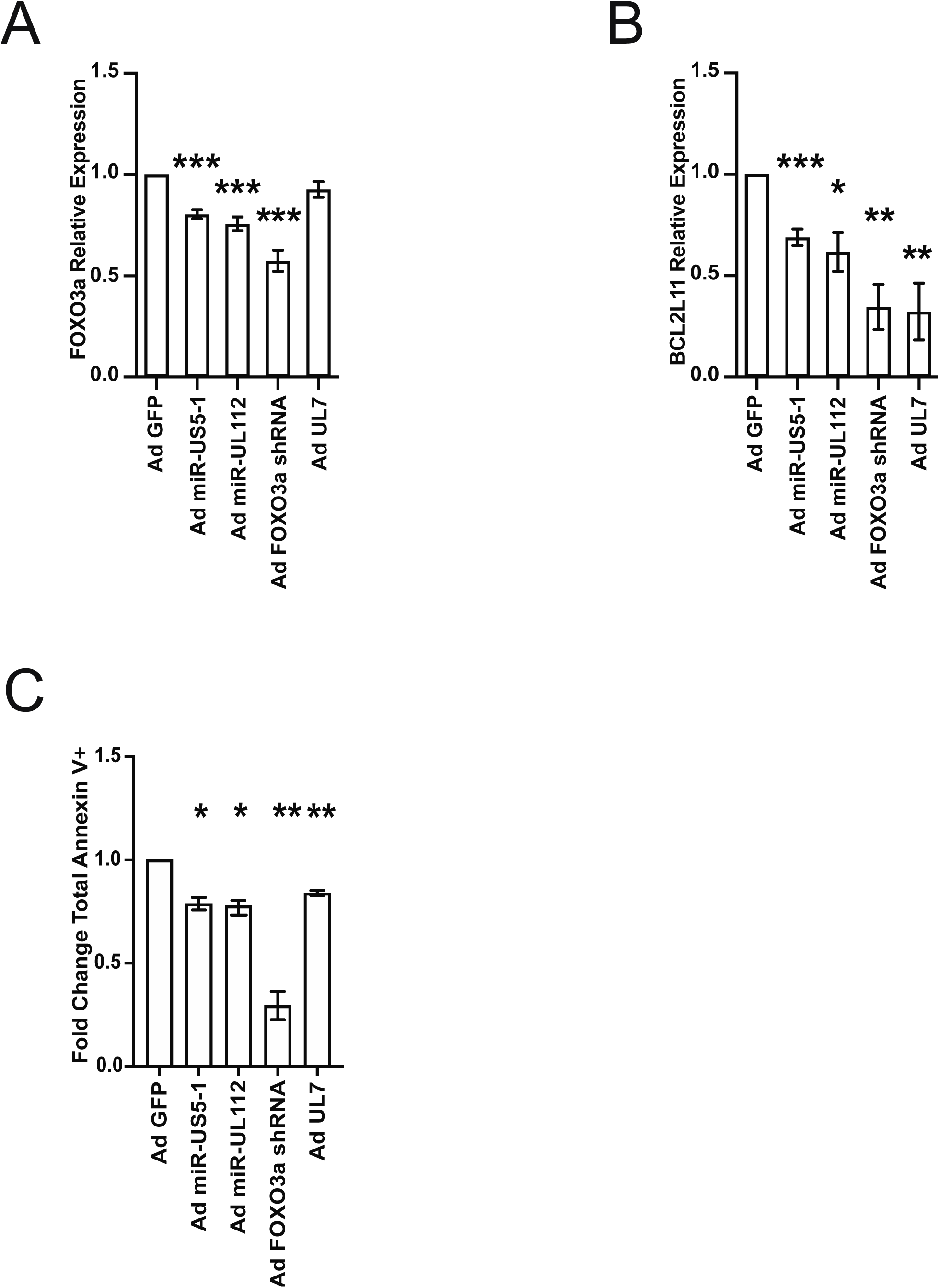
HCMV UL7, miR-US5-1 and miR-UL112-3p protect CD34^+^ HPCs from apoptosis. CD34^+^ HPCs were transduced at an MOI of 500 with Ad GFP, Ad miR-US5-1, Ad miR-UL112-3p, Ad FOXO3a shRNA, or Ad UL7 for 48h and then FACS-isolated for viable CD34^+^ GFP^+^ HPCs. RNA was isolated using Trizol and qRT-PCR for *FOXO3a* (A) and *BCL2L11* (B) was performed. Values are means±standard error of the means (SEM) (error bars) compared to Ad GFP for three independent CD34^+^ donors. Statistical significance was determined by unpaired Student’s t-test (*, *P<0*.*05*, **; *P<0*.*005*; ***; *P<0*.*0005*). C) CD34+ HPCs were transduced as above and analyzed by flow cytometry for apoptosis induction after 72h. The total population was gated on total single, CD34^+^, GFP^+^ HPCs and analyzed for Annexin V^+^ (early apoptotic cells) plus Annexin V^+^ and Viability Dye^+^(late apoptotic and dead cells). The fold change in total Annexin V^+^ cells compared to the Ad GFP group for three independent donors is shown. Statistical significance was determined by paired Student’s t-test (*, *P<0*.*05*, **; *P<0*.*01*).

### UL7, miR-US5-1 and miR-UL112-3p decreases *BCL2L11* gene expression in HCMV-infected CD34^+^ HPCs

FOXO3a is mostly found in the nucleus of primitive hematopoietic cells and progenitor cells, as shown in Figure 2C. We hypothesized that UL7, miR-US5-1 and miR-UL112-3p could act together to reduce FOXO3a levels and activity at early times post-infection to block the induction of the pro-apoptotic transcript *BCL2L11*. As shown in Figure 5, UL7 is expressed at 2 days post wild-type HCMV infection in four independent donors. The expression of HCMV miR-US5-1, and miR-UL112-3p in CD34^+^ HPCs at the same time point has previously been reported (39). After sorting for a pure population of GFP^+^, CD34^+^ HPCs at 2dpi we observed that FOXO3a transcript was significantly increased in TB40EΔmiR-US5-1+UL112-3p compared to wild-type infection (p=0.0003, Figure 5B), while *BCL2L11* mRNA levels were higher in both TB40EΔmiR-US5-1+UL112-3p (p<0.0001) and TB40EΔUL7 (p=0.008) (Figure 5C). These results are consistent with the hypothesis that the virus is inactivating FOXO3a using both viral miRNAs and a viral protein to prevent expression of the pro-apoptotic gene *BCL2L11* and to promote survival of HCMV-infected CD34^+^ HPCs.

**Figure 5.**
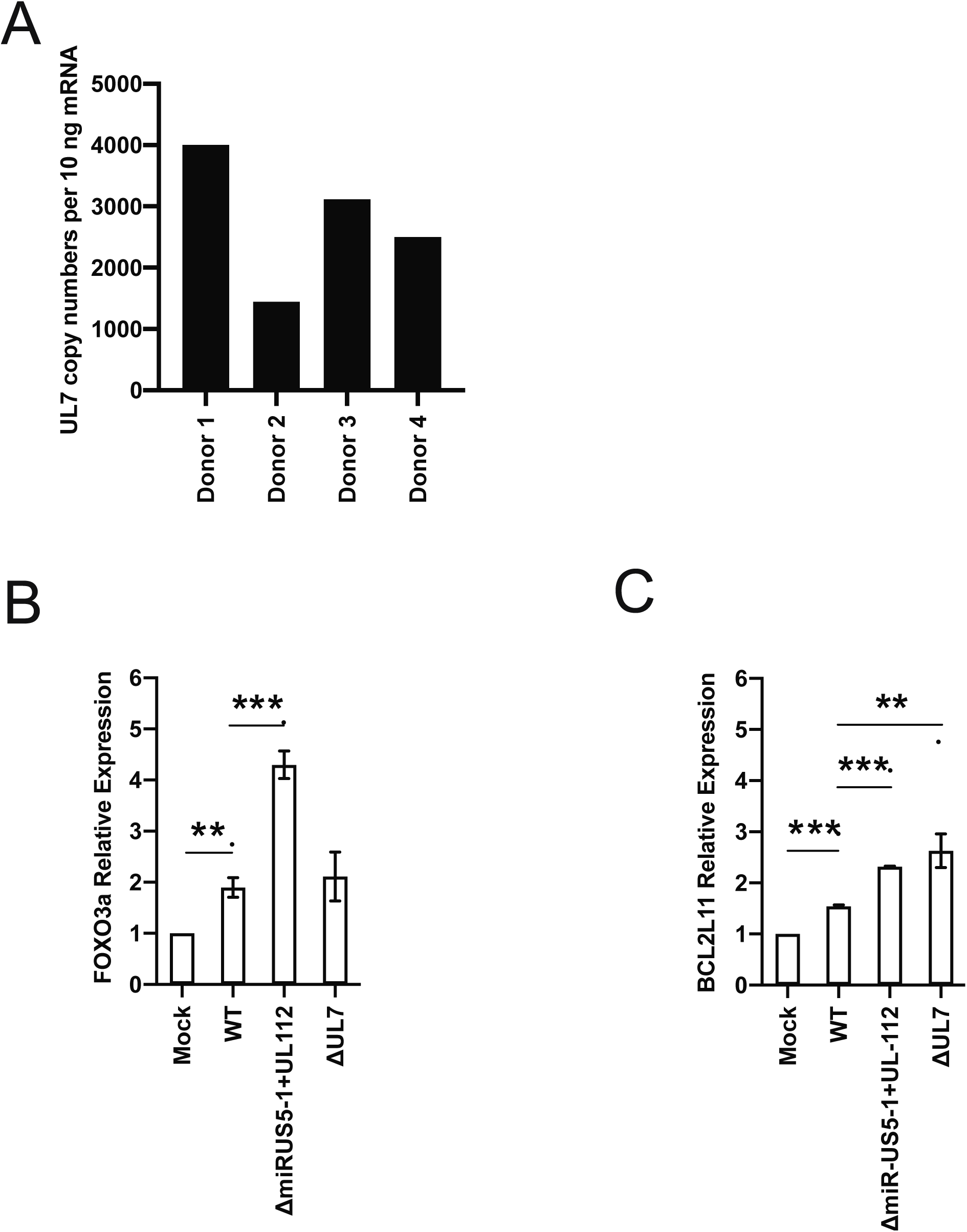
Effect of UL7, miR-US5-1 and UL112-3p deletion on FOXO3a and BCL2L11 expression in HCMV infected CD34^+^ HPCs. A) CD34^+^ HPCs were infected with WT TB40E for 48 hours, and then FACS-isolated for viable CD34^+^ GFP^+^ HPCs. RNA was isolated using Trizol and qRT-PCR for UL7 was performed. B) CD34^+^ HPCs were infected with WT TB40E, ΔmiR-US5-1+UL112-3p, or ΔUL7 for 48 hours, and then FACS-isolated for viable CD34^+^ GFP^+^ HPCs. RNA was isolated using Trizol and qRT-PCR for *FOXO3a* (B) and *BCL2L11* (C) was performed. Values are means±standard error of the means (SEM) (error bars). Statistical significance was determined by unpaired Student’s t-test (n=4 *, *P<0*.*05*, **; *P<0*.*005*; ***; *P<0*.*0005*).

## Discussion

In the present study, we report a novel observation whereby HCMV pUL7 together with miR-US5-1 and miR-UL112-3p mediate the downregulation and inactivation of the transcription factor FOXO3a prior to establishment of latency in CD34^+^ HPCs. These data implicate multiple HCMV gene products in regulation of FOXO3a, suggesting that modulation of FOXO3a function is critical during HCMV infection of CD34^+^ HPCs. Indeed, we found that inactivation of FOXO3a leads to downregulation of the pro-apoptotic gene *BCL2L11* and protection from apoptosis.

Jonsson *et al*. previously reported that Flt-3L induces AKT/PKB phosphorylation, which inactivates FOXO3a leading to the survival of a murine progenitor cell line stably expressing Flt-3R (FDC-P1/flt3) (33). We also observed that Flt-3L promotes FOXO3 phosphorylation via AKT signaling (Figure 1A, B) and inactivation of the transcription factor (Figure 1C) in telomerized human fibroblasts over-expressing the Flt-3R. Interestingly, we also observed that pUL7 promotes a time-dependent phosphorylation of FOXO3a via the Flt-3R (Figure 1A and 1B) and nuclear to cytoplasm translocation (Figure 1E), however the preferential intracellular signaling pathway stimulated by pUL7 is ERK/MAPK (Figure 1C). Specifically, pUL7 stimulation induces phosphorylation of FOXO3a on Ser294 (Figure 1D, 2A), which along with Ser344 and Ser425 are residues of FOXO3a that are phosphorylated by ERK. ERK-mediated phosphorylation induces FOXO3a degradation via the murine double minute-2 (MDM2) signaling pathway, which consequently stimulates cell survival and anti-apoptotic gene expression (34). Indeed, we demonstrated that phosphorylation of FOXO3a by pUL7 leads to nuclear exclusion in CD34^+^ HPCs (Figure 2C) and downregulation of the pro-apoptotic gene *BCL2L11* (Figure 2D). Besides the regulation of FOXO3a through phosphorylation, we found that HCMV encodes two miRNAs that are capable of downregulating endogenous FOXO3a mRNA and protein. Indeed, viral miRNAs regulate numerous cellular and viral processes critical for efficient viral replication including immune evasion, proinflammatory cytokine production and signaling, cell survival and virion assembly compartment formation (reviewed in (8)). We previously published that HCMV miR-US5-1 and miR-UL112-3p target the IкB kinase (IKK) complex components IKKα and IKKβ to limit production of proinflammatory cytokines in response to interleukin 1b (IL-1β) and tumor necrosis factor alpha (TNFα) (40). In this study, we show that the same two HCMV miRNAs target the 3’ UTR of FOXO3a (Figure 3A), downregulating endogenous RNA and protein (Figure 3B-F). In hematopoietic cells, cellular miRNAs from the miR-212/132 cluster regulate FOXO3a mRNA and protein levels. Both deletion and overexpression of this miRNA cluster results in altered hematopoiesis during aging, suggesting that precise FOXO3a regulation via miRNA expression in the bone marrow is a critical component of hematopoiesis (41).

An understanding of the role FOXO transcriptional factors in the biology of herpesviruses remains incomplete. We know from the literature that Epstein-Barr virus (EBV) latent membrane protein 1 inactivates FOXO3a via the PI3K/AKT pathway, leading to expression of miR-21 (42) and upregulation of MCL-1, both of which reduce apoptosis (43). Similarly, the LANA2 protein from Kaposi’s sarcoma-associated herpesvirus (KSHV) functionally interacts with FOXO3a and inhibits the transactivation of *BCL2L11* promoter (44). Chemical inhibition or knockdown of the related FOXO1 protein has been shown to increase the intracellular ROS level that are sufficient to disrupt KSHV latency and promote viral lytic reactivation (45). A recent publication by Hale *et al*. (46) showed an important role for FOXO proteins in HCMV reactivation from latency. FOXO1 and FOXO3a transcription factor activity is essential for inducing HCMV reactivation by activating the alternative major immediate early promoters iP1 and iP2. This study builds on the observation that IE protein expression following reactivation stimuli is dependent not on expression from the MIEP, but predominantly from the novel internal promoters (46, 47). Thus, it appears that herpesviruses utilize multiple different mechanisms to regulate FOXO protein function to persist in infected cells and in some instances to promote viral reactivation. Our data are similar to what has been reported for KHSV LANA2 protein (44). Indeed we found that UL7, miR-US5-1, and miR-UL112-3p inactivation of FOXO3a is important to promote survival of CD34^+^ HPCs when they are independently expressed (Figure 4C) as well as in the context of the viral infection, as demonstrated by the increased expression of *BCL2L11* during infection with the HCMVΔmiR-US5-1+ UL112-3p and ΔUL7 mutant viruses (Figure 5D).

HCMV is known to promote an anti-apoptotic response through the activation of signaling pathways in order to ensure long-term survival in non-permissive cells (reviewed in (17)). Indeed, the protective phenotype upon entry in CD34^+^ HPCs is achieved by virus induced ERK-MAPK signaling that leads to the degradation of pro-apoptotic proteins PUMA and BIM, alongside with an up-regulation of MCL-1 levels (20, 21). Our findings show that the ERK-MAPK pathway is important for the UL7-mediated phosphorylation and inactivation of FOXO3a, resulting in downregulation of BIM transcript and protein and ultimately in protection from apoptosis (Figure 4C). The role of HCMV miRNAs in protecting non-permissive cells from apoptosis is unknown. Here, we report that HCMV miR-US5-1 and miR-UL112-3p protect CD34^+^ HPCs from apoptosis by targeting FOXO3a and ultimately decreasing *BCL2L11* gene expression. Among herpesviruses, EBV and KHSV are known to use viral miRNAs as a tool to avoid apoptosis both at early stages of infection and upon cell transformation. EBV-miR-BART5-5p regulates pro-apoptotic PUMA, while ebv-miR-BART-cluster 1 and ebv-miR-BART12 target BIM (48, 49). BCL2-associated death promoter (BAD) protein is downregulated by ebv-miR-BART20-5p (50). Finally, pro-apoptotic protein caspase 3 was identified as being a target of ebv-miR-BART16 and ebv-miR-BART1-3p (51). KHSV encodes three viral miRNAs, kshv-miR-K12-1, kshv-miR-K12-3, and kshv-miR-K12-4-3p, that cooperatively suppress caspase 3 expression and reduce apoptosis in infected cells (52).

This study illustrates for the first time a synergy between a viral gene and two miRNAs in regulating a cellular target, FOXO3a, to promote CD34^+^ HPC survival. Greater understanding of how UL7, miR-US5-1, and miR-UL112-3p work together in the context of viral infection is important to better define their role in establishment of HCMV latency and viral reactivation.

## Materials and Methods

### Cells, reagents and viruses

CD34^+^ HPCs were isolated from fetal liver tissue obtained from Advanced Bioscience Resources. The tissue was manually disrupted then digested with DNase, Collagenase and Hyaluronidase, and CD34^+^ HPCs were isolated using magnetic beads (Miltenyi Biotech) as previously described (53). Cells for HCMV infection were cultured in IMDM (HyClone) supplemented with 10% BIT 9500 serum substitute (StemCell Technologies), 50 μM 2-mercaptoethanol, 2nM L-glutamine, 20 ng/ml human low-density lipoprotein (MilliporeSigma) as previously described (54). Human bone marrow CD34^+^ HPCs were obtained from Stem cell Technologies and recovered in SFEMII (Stem Cell Technologies) supplemented with 10% BIT 9500 serum substitute. RS4;11 (bone marrow lymphoblast) cells were obtained from the American Type Culture Collection. RS4;11 cells were grown in RPMI-1640 (HyClone) medium supplemented with 10% fetal bovine serum (FBS) (HyClone), 4.5 g/l glucose, L-glutamine and sodium pyruvate, and antibiotics [penicillin (10 units/ml)–streptomycin (10 μg/ml)]. Telomerized human fibroblasts (THF) were a gift from Dr Victor DeFilippis (OHSU). THF cells were cultured with Dulbecco’s modified Eagle’s medium (DMEM) (Cellgro) supplemented with 10% FBS, penicillin, streptomycin, and L-glutamine. Normal human dermal fibroblasts (NHDF) were cultured in conditions identical to the THF cells. Human Embryonic Kidney 293 cell line (HEK293) (Microbix Biosystems Inc) were cultured in Minimum Essential Medium (MEM) (Cellgro) supplemented with 10% FBS, penicillin, streptomycin, and L-glutamine. Stably transduced THF expressing human Flt-3R were generated by transduction with the lentivirus pLenti-FLT3-mGFP-P2A-Puro (OriGene Technologies RC211459L4V) followed by GFP purification after one week of Puro (800 μg/ml) selection. Recombinant UL7 protein from TB40E (TB40E: EF999921; UL7 protein: ABV71537.1) was generated at RayBiotech as previously described (31). Recombinant mouse IgG Fc and recombinant human Flt-3L were purchased at R&D systems. Quizartinib (AC220), PD98059, LY294002, LY303511 inhibitors were purchased at Selleckchem and resuspended in DMSO. HCMV TB40, HCMV TB40ΔUL7, HCMV TB40ΔmiR-US5-1ΔmiR-UL112-3p stocks and titers were generated as previously described (31, 40).

### Immunofluorescence

CD34^+^ HPCs from 3 independent donors were thawed and plated in StemSpan SFEM II media (Stemcell Technologies) containing 10% BIT serum substitute, in a 5% CO2 humidified incubator and recovered overnight (15-18h) at 37oC. A total of 50,000 cells were seeded per well in an 8-well chamber slide (Thermo Scientific) in fresh SFEM II media without additional supplements. The HPCs were then either unstimulated (mock), or stimulated for 6 h with 50 ng/mL of recombinant human Flt-3L or 50 ng/mL of recombinant UL7 protein. HPCs were fixed in 4% methanol-free formaldehyde for 15 min, then washed 3 times in PBS. After permeabilizing in PBS containing 0.1% Triton X-100 for 10 min and washing in PBS, the HPCs were blocked for 1 h in freshly prepared 5% normal goat serum with 0.3% Triton X-100 in PBS. After washing in PBS, the HPCs were incubated overnight at 4°C in the primary antibodies rabbit-anti-FoxO3a (Cell Signaling), rabbit-anti-p27 ^kip1^ (Abcam), or mouse-anti-CD34 (Biolegend), all diluted 1:100 in antibody dilution buffer (PBS containing 1% bovine serum albumin (BSA) and 0.3% Triton X-100). The HPCs were washed twice in PBS, then incubated 1 h in the secondary antibody goat-anti-rabbit AF594 (Invitrogen) and goat-anti-mouse AF488 (Invitrogen) diluted at 1:1,000 in antibody dilution buffer. The HPCs were mounted under a coverslip with Dapi-Fluoromount-G clear mounting media (SouthernBiotech) and imaged using an EVOS-FL fluorescent microscope with a 100X objective. The image analysis was performed using ImageJ.

### Adenoviruses

HCMV UL7 containing an HiBiT sequence (5’-gtgagcggctggcggctgttcaagaagattagc-3’) after the signal peptide (nucleotide 142) or the regions approximately 100bp up and downstream of miR-US5-1 and miR-UL112 were cloned into the shuttle vector pAdTrack-CMV and transformed into TOP10 cells. After a 24h incubation at 37°C under kanamycin selection, colonies were selected and screened for the insert by restriction enzyme digest using *KpnI* and *XhoI*. The pAdTrack plasmids were then linearized by digesting with restriction endonuclease *Pme I*, and subsequently recombined into *E. coli* BJ5183 cells containing the adenoviral backbone plasmid pAdEasy-1(AdEasier-1 cells). pAdTrack-CMV (Addgene plasmid #16405) and AdEasier-1 cells (Addgene #16399) were a gift from Bert Vogelstein (He et al, 1998). Recombinants were selected for kanamycin resistance, and the recombination confirmed by restriction endonuclease analyses. Finally, the recombinant plasmids were linearized with *PacI* before transfection with Lipofectamine 2000 (ThermoFisher) into the adenovirus packaging cell lines HEK293 cells. The control vector Ad GFP, Ad miR-UL112-3p, Ad miR-US5-1, and Ad UL7 were produced, purified and titered in HEK293, as previously described (27). The Ad-GFP-U6-h-FOXO3a-shRNA (shADV-209273) was purchased at Vector Biosystems, Inc.

### qRT-PCR

Total RNA was isolated from infected, transfected or treated cells using the Trizol RNA isolation method following the manufacturer’s directions. cDNA was prepared using 10 to 1000 ng of total RNA and random hexamer primers. Samples were incubated at 16°C for 30 minutes, 42°C for 30 minutes and 85°C for 5 minutes. Real-time PCR (Taqman) was used to analyze cDNA levels in transfected or infected samples. An ABI StepOnePlus Real Time PCR machine was used with the following program for 40 cycles: 95°C for 15 sec and 60°C for one minute. FOXO3a, BCL2L11 and 18S primer/probe sets were obtained from ThermoFisher Scientific. Relative expression was determined using the ΔΔCt method using 18S as the standard control. For UL7 expression we used the following set of primers and probe: UL7_F primer 5’-ACTACGTGTCGTCGCTGGATT-3’; UL7_R primer 5’-ACAACTTCCACCACCCCATAAT; UL7 probe 6FAM-CATGGCCTTGGTAGGTG-MGBNFQ. qRT-PCR assays for miR-US5-1 and miR-UL112 were purchased from ThermoFisher Scientific.

### Luciferase Reporter Assay

The putative 3’ UTR of FOXO3a was cloned into the dual luciferase reporter pSiCheck2 (Clontech) using the following primers: FOXO3a F: GGCAAGGCAGCACAAAACAG, FOXO3a R: GCTTTATTTACATGCGTCACC. Site-directed mutagenesis was performed using the QuikChange PCR method. To mutate the potential miR-US5-1 site within the FOXO3a 3’ UTR the following primers were used: FOXO3a SDM 5-1 F: CATTTTAAAAATTCAGAACTCCTGTTAATGGGAGG, FOXO3a SDM 5-1 R: CCTCCCATTAACAGGAGTTCTGAATTTTTAAAATG. To mutate the potential miR-UL112 site within the FOXO3a 3’ UTR the following primers were used: FOXO3a SDM 112 F: CACATTTTAAAAATTCAGAACTCCTGTTAATGGGAGGATC, FOXO3a SDM 112 R: GATCCTCCCATTAACAGGAGTTCTGAATTTTTAAAATGTG. 293T cells seeded into 96-well plates were co-transfected in triplicate with 100ng of plasmid and 100fmol of miRNA mimic (custom designed; IDT) using Lipofectamine 2000 (Invitrogen). Cells were incubated overnight, and then harvested for luciferase assay using the Dual-Glo Reporter Assay Kit (Promega) according to the manufacturer’s protocol. Luminescence was detected using a Veritas Microplate Luminometer (Turner Biosystems).

### Immunoblotting

Cytoplasmic and nuclear extracts were obtained using the NE-PER Nuclear and Cytoplasmic Extraction kit (Pierce Biotechnology). Total protein extracts were prepared in cell lysis buffer [20 mM Tris-HCl pH7.5, 150 mM sodium chloride (NaCl), 1%(v/v)polyethylene glycol octyl phenol ether (TritonX-100), 2.5mM sodium pyrophosphate,1mMethylenediaminetetraaceticacid (EDTA),1% (w/v) sodiumorthovanadate, 0.5 μg/ml leupeptin,1mM phenyl-methanesulfonyl fluoride (PMSF)] and run on an 8-12% SDS-PAGE, transferred to Immobilon-P Transfer Membranes (Milipore Corp., Bedford, MA), and visualized with antibodies specific for P-Flt3R (Tyr591, 54H1 Cell Signaling), Flt-3R (8F2, Cell Signaling), FOXO3a (Cell Signaling), P-FOXO3a (Ser253, Cell Signaling), P-FOXO3a (Ser294, Cell Signaling), P-Akt (Thr308, Cell Signaling), Akt (C73H10, Cell Signaling), ERK1 (C16, Santa Cruz), P-ERK1/2 (Thr202/Tyr204, Cell Signaling), PARP (Cell Signaling), P-6S Ribosomal protein (Ser235/236, D57.2.2E Cell Signaling), S6 Ribosomal protein (5G10, Cell Signaling), BIM (C34C5, Cell Signaling), GAPDH (Abcam). Relative intensity of bands detected by western blotting was quantitated using ImageJ software.

### Adenovirus transduction of CD34^+^ HPCs

CD34^+^ HPCs were freshly isolated or thawed and recovered for 3 hours in IMDM containing 1%FBS, 1%PSG, and stem cell cytokines (SCF, FLT3L, IL-3, and IL-6). Following recovery, cells were washed in PBS, and resuspended at low volume in IMDM containing 10% BIT serum supplement, L-glutamine, low density lipoproteins, B-ME, and stem cell cytokines as previously described (54) in a low binding 24-well plate (Corning low-attachment hydrocell). HPCs were infected with adenoviruses at an MOI of 500 for 4 hours with continual rocking then spin infected at 300g for 30min, resuspended, and cultured overnight. Culture conditions were supplemented with additional media and infection continued for a total of 48-72 hours. Samples were then FACS isolated for pure populations of transduced HPCs (viable, CD34^+^, GFP^+^) as previously described (31), or analyzed by flow cytometry as described below.

### Flow Cytometry Analysis for Apoptosis

CD34^+^ HPCs were transduced with adenoviruses for 72 hours were washed in PBS and stained with Fixable Viability Dye eFluor 780 (Initrogen/eBioscience). Cells were washed twice in FACS buffer, blocked, and stained with surface antibodies for stem cell markers including CD34 as previously described (53). HPCs were stained with Annexin V (eBioscience) according to the manufacturer’s instructions and then washed fixed with 2% formalin as previously described (53). Data was acquired on an LSRII flow cytometer (BectonDickson) running FACSDIva software and data was analyzed with FlowJo v10.7 (TreeStar).

### Statistical Analysis

Data are expressed as the mean ± s.e.m. Statistical analysis was performed using GraphPad Prism (v8) for comparison between groups using Student’s t-test as indicated. A value of p<0.05 or lower was considered significant and exact p values are indicated.

## Acknowledgements

This work was supported by grant P01 AI127335 from the National Institute of Allergy and Infectious Diseases, NIH funded to P.C. M.H.H is supported by National Institute of Health R37 AI21640 and P01 AI127335. The funder had no role in study design, data collection and analysis, decision to publish, or preparation of the manuscript.

We thank Andrew Townsend for graphics assistance. We gratefully acknowledge Jay Nelson, Daniel Streblow, Felicia Goodrum, and Andrew Yurochko for helpful discussions.

